# Emergence of diauxie as an optimal growth strategy under resource allocation constraints in cellular metabolism

**DOI:** 10.1101/2020.07.15.204420

**Authors:** Pierre Salvy, Vassily Hatzimanikatis

## Abstract

The sequential rather than simultaneous consumption of carbohydrates in bacteria such as *E. coli*, a phenomenon termed diauxie, has been hypothesized to be an evolutionary strategy which allows the organism to maximize its instantaneous specific growth, thus giving the bacterium a competitive advantage. Currently, computational techniques used in industrial biotechnology fall short of explaining the intracellular dynamics underlying diauxic behavior, in particular at a proteome level.

Some hypotheses postulate that diauxie is due to limitations in the catalytic capacity of bacterial cells. We developed a robust iterative dynamic method based on expression- and thermodynamically enabled flux models (dETFL) to simulate the temporal evolution of carbohydrate consumption and cellular growth. The dETFL method couples gene expression and metabolic networks at the genome scale, and successfully predicts the preferential uptake of glucose over lactose in *E. coli* cultures grown on a mixture of carbohydrates. The observed diauxic behavior in the simulated cellular states suggests that the observed diauxic behavior is supported by a switch in the content of the proteome in response to fluctuations in the availability of extracellular carbon sources. We are able to model both the proteome allocation and the proteomic switch latency induced by different types of cultures.

Our models suggest that the diauxic behavior of the cell is the result of the evolutionary objective of maximization of the specific growth of the cell. We propose that genetic regulatory networks, such as the *lac* operon in *E. coli*, are the biological implementation of a robust control system to ensure optimal growth.

## Introduction

In his pioneering work on the growth of bacterial cultures, the French biologist Jacques Monod [1] observed that the growth of *Escherischa coli* (*E. coli*) in a mixture of carbohydrates followed two distinct exponential curves separated by a plateau — a phenomenon he called diauxie. Hypothesized to allow optimal growth of the culture [2], this cellular behavior corresponds to the sequential consumption of sugars, where one sugar is preferentially consumed, and the second is consumed after depletion of the first Although current optimality-based computational models can predict diauxie, these lack a detailed description of protein dynamics during the phenomenon [3]. Diauxie is an evolved, complex behavior, and its occurrence is controlled by the regulation network of the *lac* operon in *E. coli* [4]. The emergence of such a control mechanism is the product of evolutionary pressure, and being able to fully elucidate its raison d’être in terms of cell physiology is an important milestone to understand and better engineer the intracellular dynamics of bacterial growth. There is thus a need for a formulation describing diauxie at the proteome level.

Genome-scale models of metabolism (GEMs) combine constraint-based modeling and optimization techniques to study cell cultures [5–7]. A key method for studying GEMs is flux balance analysis (FBA) [8], which formulates a linear optimization problem that employs stochiometric constraints through the mass conservation of metabolites given their synthesis and degradation reactions. Under the typical steady-state and growth-rate maximization assumptions, FBA models predict the simultaneous consumption of two or more carbon sources to achieve the maximum possible growth [3]. However, this contradicts Monod’s observation of distinct, sequential phases of carbon consumption and suggests that diauxie does not come from stoichiometric constraints.

To account for diauxie beyond stoichiometric modeling, we looked into other biological features. Because a cell has a physiological constraint on the total amount of enzymes it can house, which we will call a proteome allocation constraint, it is likely that the cell will preferentially distribute its limited catalytic capacity towards pathways that utilize the most efficient substrate/enzyme combination [2, 9]. Therefore, models that account for proteome limitation in cells may be able to account for diauxie. Towards this end, the role of protein limitation in diauxie was demonstrated by Beg *et al.* [10] with their formulation of FBA with molecular crowding (FBAwMC). Their method correctly predicts the uptake order of 5 different carbon sources in a batch reactor, using a proteome allocation constraint. In a push towards more global models, models of metabolism and expression (ME-models) [11, 12] include proteome allocation, but also gene expression mechanisms, a modeling paradigm that is ideal for studying diauxie at the proteome level. ME-models also fully describe the requirements of enzyme synthesis, degradation, and dilution effects, as well as mRNA and enzyme concentrations.

Since the diauxie is also a time-dependent phenomenon, we chose to complement ME-models with a dynamic modeling approach. Dynamic FBA (dFBA) [13] is a generalization of FBA for modeling cell cultures in time-dependent environments. In its original static optimization approach (SOA) formulation, the time is discretized into time steps, and an FBA problem is solved at each step. At each iteration, kinetic laws and the FBA solution are used to update the boundary fluxes, extracellular concentrations, and cell concentration, based on the amount of substrate consumed, byproducts secreted, and biomass produced by the cells. We expected that the combination of a dFBA and ME-models would yield a formulation that can describe diauxie at the proteome level.

However, we identified three major challenges in the conception of dynamic models of metabolism and expression. First, while dFBA studies of metabolic networks can be solved by common linear solvers, ME-models are non-linear by nature, and significantly more complex. The new species and reactions introduced and considerations of the interactions between enzyme expression and metabolism result in nonlinear problems that are often 1–2 orders of magnitude bigger in terms of constraints and variables than the corresponding linear (d)FBA problem. The increase in complexity is compounded when iteratively solving an optimization problem. As a result, combining ME-models and dynamic studies brings along difficulties that arise from the high computational cost of solving multiple times, with different conditions, these large, non-linear problems. Second, the use of iterative methods presents the additional challenge of alternative solutions, which can span several physiologies. It is thus necessary to find, for each time step, a suitable representative solution that will be used to integrate the system. This also poses the problem of finding a set of initial conditions for the system. Third, the current state-of-the-art models present limitations at the proteome level. Lloyd *et al.* [14] developed an efficient ME-model for *E. coli*, and Yang *et al.* used it to formulate a dynamic analysis framework (dynamicME) [15] similar to dFBA. However, the assumptions introduced to alleviate the computational complexity of their model limit some aspects of the modeling capabilities of their method (Supplementary Note S1). In particular, DynamicME forces a strict coupling between enzyme concentrations and fluxes. However, a change in the growth conditions will trigger a change in the proteome allocation to adapt to a new metabolic state, or lag phase. During that time, it is expected that some previously active enzymes will not be able to carry flux in the new conditions. Therefore, enzyme flux and concentration will decouple, unless the enzyme composition of the proteome changes at the same rate as the environment. As a result, the method cannot simulate lag phase during glucose depletion and proteome reallocation.

Both dynamic models and models including gene expression mechanisms are important components in the development of successful predictive biology [16]. We propose a dynamic method which tackles the challenges mentioned above and models diauxie at the proteome level. To this effect, we used our recently published framework for ME-models, ETFL [17]. The formulation of ETFL permits the inclusion of thermodynamics constraints in expression models, as well as the ability to describe the growth-dependent allocation of resources. ETFL is faster than previous ME-model formulations, thanks to the use of standard mixed-integer linear programming (MILP) solvers [17]. We herein leverage ETFL for dynamic analysis, in a method called dETFL. It includes a method based on Chebyshev centering to robustly select a representative solution from the feasible space at each time step. The representative solution captures phenotypic and genotypic differences between cells precultured in different media. (d)ETFL solves the problem of computational accuracy lacking in previous models by performing a systematic scaling of its constraints, eliminating the need for dedicated solvers. This allows models to be solved efficiently, without resorting to a strict coupling of enzymes and fluxes. As a result, whole-proteome reconfiguration during sugar consumption can be simulated, which will enable the modeling of the lag phase in diauxie.

Herein we model the emergence and dynamics of diauxie arising at the proteome level. We first propose a small conceptual model of a cell, with a limited in proteome, and demonstrate its ability to predict diauxie under a minimal set of assumptions. Using the dETFL method, we subsequently show these assumptions hold in *E. coli*, and reproduce experimental results of bacterial growth. Finally, we apply the dETFL framework to the growth of *E. coli* in a glucose/lactose mixture in a batch reactor, and demonstrate that it robustly predicts diauxie as well as the preferential consumption of glucose over lactose. Overall, dETFL offers a method to robustly survey intracellular dynamics of cellular physiology under changing environmental conditions.

## Results

### Conceptual model for the emergence of the diauxie phenotype from proteome limitation

We designed a simplified conceptual model, to illustrate diauxie from proteome limitations, as described in Fig.1-a. The model includes both glucose and lactose as substrates, and it is a simplified version of the *E. coli* metabolism based on four considerations:

**Fig 1.**
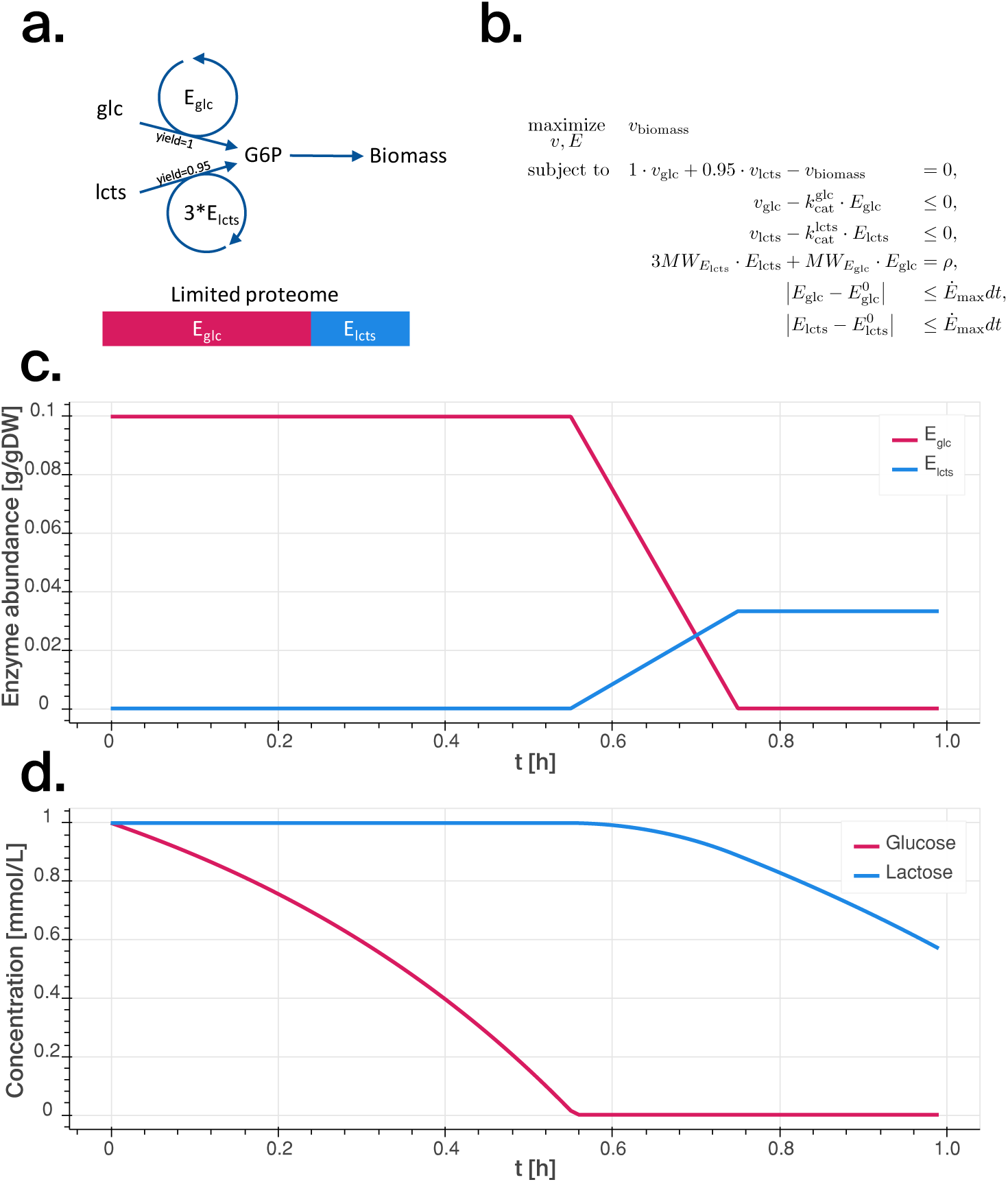
**a.** Conceptual model used for the preliminary analysis, where “glc” stands for glucose, “lcts” for lactose. The catalytic efficiency of the enzymes are assumed to be the same. Three enzymes are assumed to be necessary to produce the intermediate metabolite G6P from lactose, and only one enzyme is required from glucose. **b.** Optimization problem used to represent the model. *v* are fluxes, *E* are enzyme concentrations, *E*^0^ are reference values, *MW* are molecular weights, *ρ* is the mass fraction of the cell occupied by the enzymes we consider, 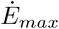 is the maximal variation of enzyme concentration over time, and *dt* is the integration interval. **c.** Enzyme content over time for the conceptual model growing on a mixed substrates. **d.** Changes in sugar content of the batch reactor over time.

(C1) The biomass carbon yield on glucose is slightly higher than that of lactose [18]
(C2) Glucose and lactose are taken up and converge to a common intermediate metabolite, glucose 6-phosphate (G6P). Glucose is transformed into G6P by a glucokinase. The lactose pathway (Leloir pathway) splits the lactose, a disaccharide, into its glucose and galactose subunits. The galactose is then converted to G6P by a series of enzymes.
(C3) The Leloir pathway requires one enzyme to split the lactose into glucose and galactose, four enzymes to convert galactose into glucose-1-phosphate [19, 20], and one to convert glucose-1-phosphate into G6P; this bring the total to six enzymes needed for the synthesis of two G6P, which is equivalent to three enzymes per G6P;
(C4) The molecular weight of the each each of the enzymes in the lactose pathway is around 60-90 kDa [21], which is heavier than the 33 kDa glucokinase (Uniprot ID A7ZPJ8)).

Based on these considerations, we devised a conceptual model of glucose and lactose metabolism for *E. coli*. The model accounts for the consumption of the two substrates, which both synthesize an intermediate metabolite that is then used to make biomass. We thus made five modeling assumptions:

(A1) Glucose has a slightly higher carbon yield than lactose — based on (C1).
(A2) The glucose and lactose metabolism leading to the intermediate G6P are catalyzed by two different enzymes — based on (C2).
(A3) The molecular weights of the enzymes are the same, and three times more enzymes are required for lactose metabolism than for glucose metabolism — based on (C3).
(A4) The catalytic activities of the two enzymes synthesizing G6P are similar.
(A5) The variation of enzyme concentrations reaches a maximum at each time step.
(A6) The total enzyme amount in the cell is limited.

The mathematical formulation of the problem (Fig.1-b) involves one mass balance, one conservation equation of the total enzymes, two inequalities that constrain the metabolism for glucose and lactose as a function of the corresponding enzymes concentrations, and two enzyme variation constraints. Due to total enzyme conservation, the two maximum activity constraints are not independent. This constraint is similar to that found in other approaches accounting for proteome allocation such as FBAwMC [10].

The conceptual model is able to predict diauxic behavior in our system. The model shows the preferential consumption of glucose over lactose (Fig.1-d), controlled by a switch in the proteome composition over time (Fig.1-c). The diauxic phenomenon is due the fact that the system will invest all the (limited) enzyme resources into the metabolism of glucose, which is both the highest yielding substrate ((C1), (A1)), and the one with least enzyme requirements ((C3) and (A3)). As the glucose is depleted, the uptake flux is reduced and the system gradually allocates part of its proteome for enzymes needed for lactose metabolism. This gradual proteome reallocation corresponds to the observed lag phase in an experimental system. While this conceptual model lacks catabolite repression mechanisms, it can still describe diauxie phenomenon from only the proteome capacity constraint.

The lack of proteome limitation and enzyme catalytic constraints is why the original FBA approach fails to predict diauxie, which leads to the simultaneous utilization of both sugar substrates. However, another important constraint to accurately describe the lag phase is the limitations on the rate of change in enzyme concentrations, without which the proteome switch would occur instantaneously (see Supplementary Figure S9). A cell needs time to adapt its proteome, and the limits on the rate of change of enzyme concentrations represent the catalytic limitation of the cell to break down old enzymes and synthesize new ones better adapted to the new conditions.

The conceptual model also allows for the study of the conditions under which the system switches to lactose as a carbon source, and the identification of parameters responsible for this behavior. If we note respectively for glucose and lactose, the specific growth rates on each substrate 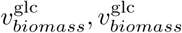, the carbon yields *Y*_glc_, *Y*_lcts_, and the catalytic rate constants 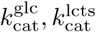 of enzymes at concentrations *E*_glc_, *E*_lcts_, then the preferred carbon source will change to lactose if and only if:

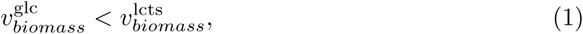

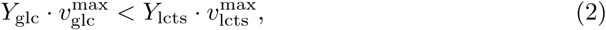

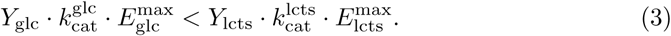

If the amount of available enzymes is represented by *ρ*, as a fraction of the total cell mass (in g.gDW^−1^), and assuming different molecular weights *MW*_*E*_, the proteome limitation constraint will be written:

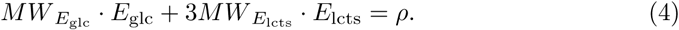

The maximal achievable values for the enzyme concentrations will be 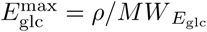 and 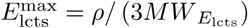. Replacing these values in Eq. 3 directly gives the condition :

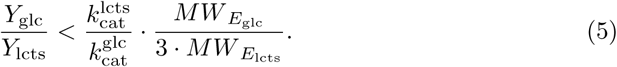

In our conceptual model, 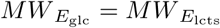, and we can simplify Eq. 5:

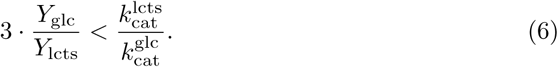

This expression identifies the boundary in the parameter space that separates the preferential use of glucose versus lactose.

These calculations can be generalized for a more realistic model, by accounting for the molecular weight of the enzymes and setting an adequate proteome fraction allocated to carbon metabolism. In practice, the catalytic efficiencies of the glycolytic enzymes are also higher than those of the Leloir pathway ((A3), see Supplementary Table S2), and the Leloir pathway enzymes are heavier ((C4), (A4)) which favors glucose consumption even more. Additionally, we did not consider the synthesis cost of the enzymes used to carry the fluxes in each pathway. Taking such property into account would also strengthen the preference towards glucose, as fewer enzymes are needed for its metabolism.

### Diauxie in genome-scale, ME-models with thermodynamic constraints

Going beyond a conceptual model, we next used dETFL to model diauxie in a ME-model of *E. coli*. This method allowed us to study metabolic switches in response to a changing environment, under the aspect for intracellular enzyme and mRNA concentrations. To do this, we studied how ME-models can describe diauxie in experiments where *E. coli* are grown in two different conditions. Firstly, we investigated the growth of *E. coli* on glucose. In this experiment, the cell exhibit overflow metabolism, or the secretion of acetate, even under aerobic conditions. Experimentally, the bacterium reutilizes the secreted acetate after glucose depletion, a form of diauxic behavior. This type of study was also used as the first proof of concept for dynamic FBA [13] Thus, we first validated the dETFL model by demonstrating its ability to model a first diauxic phenotype: overflow metabolism and acetate secretion in the presence of excess glucose, followed by acetate reutilization on glucose depletion. Secondly, we reproduced Jacques Monod’s experiment of the diauxic growth of *E. coli* in an oxygenated batch reactor [1] with a limited carbon supply made of a mixture of glucose and lactose. We aimed at reproducing the results shown in the conceptual model on a model of a real organism, and characterize the intracellular dynamics underlying the glucose/lactose diauxic behavior.

To conduct these studies, we used the *E. coli* model published by Salvy *et al.* that is based on the genome scale model by Orth *et al.* iJO1366 [22], and was assembled using ETFL. This model is significantly bigger than the conceptual model studied in the previous section, with 5295 species, 8061 reactions and 578 enzymes. A summary of the model is available in Table 1.

**Table 1.** Properties of the vETFL model generated from iJO1366.

|  |  |
| --- | --- |
| Growth upper bound $\bar{\mu}$ | $3.5h^{-1}$ |
| Number of bins $N$ | 128 |
| Resolution $\frac{\bar{\mu}}{N}$ | $0.027h^{-1}$ |
| Number of constraints | 69323 |
| Number of variables | 50010 |
| Number of species | 5295 |
| – Metabolites | 1806 |
| – Enzymes | 578 |
| – Peptides | 1433 |
| – mRNAs | 1433 |
| – tRNAs | $21 \times 2$ |
| – rRNAs | 3 |
| Number of reactions | 8061 |
| – Metabolic | 1840 |
| – Transport | 733 |
| – Exchange flux | 330 |
| – Transcription | 1433 |
| – Translation | 1433 |
| – Complexation | 578 |
| – Degradation | 2011 |
| Number of metabolites $\Delta_f G'^o$ | 1737 |
| Number of reactions $\Delta_r G'^o$ | 1787 |
| Percent of metabolites $\Delta_f G'^o$ | 93.9% |
| Percent of reactions $\Delta_r G'^o$ | 69.5% |

For the integration of the dynamic method, it is important to choose a time step that respects the quasi-steady-state assumptions on which the FBA and ETFL frameworks depend [17]. We used a time step of 0.05 h = 3 min for the numerical integration, as this is around ten times smaller than a typical doubling time for *E. coli*, and efficiently balances the integration approximation and solving time.

#### Diauxic growth on glucose and acetate

We compared the accuracy of our computational modeling of diauxie to experimental findings. Specifically, we studied the diauxic growth of *E. coli* on glucose using in batch reactors using experimental data published in Varma *et al.* [5] and Enjalbert et al. [23]. Previously, Varma *et al.* [5] used their data to validate a stoichiometric model of *E. coli* in quasi-steady state, whereas the data from Enjalbert *et al.* [23] was used to validate a population-based approach of dFBA by Succuro *et al.* [24].

To reproduce the results of these two batch growth experiments, we applied constraints to the uptake of glucose and oxygen in the dETFL model (see Materials and Methods). The initial uptake rate of glucose is set to 15 mmol.gDW^−1^.h^−1^. This value is characteristic of a typical physiology for *E. coli* growing on glucose with excess oxygen [5, 13, 22, 24].

We also matched the initial concentrations of cells, glucose, and acetate are set to the values of the experimental data. Oxygen transfer was considered free (no kinetic law on uptake) in a first approximation, as done by Succuro *et al.* [24].

Our simulations agreed with the published experimental data. The temporal evolution of the glucose and acetate concentrations in the simulated batch reactor agreed with both the Varma and Enjalbert datasets, as shown in Fig. 2-a and 2-b, respectively. The cell concentration and specific growth rate also follow a similar trend (Fig. 2-c and 2-d). Both of the simulations predict a first phase where the bacteria grow steadily on glucose, which is sustained until glucose is depleted in the medium. During that time, acetate is steadily secreted by the cell, due the overflow metabolism. When extracellular glucose is depleted, the residual acetate is consumed by the cell. We observe a sharp drop in the cell growth rate, and the simulation ends when no acetate is left in the medium.

**Fig 2.**
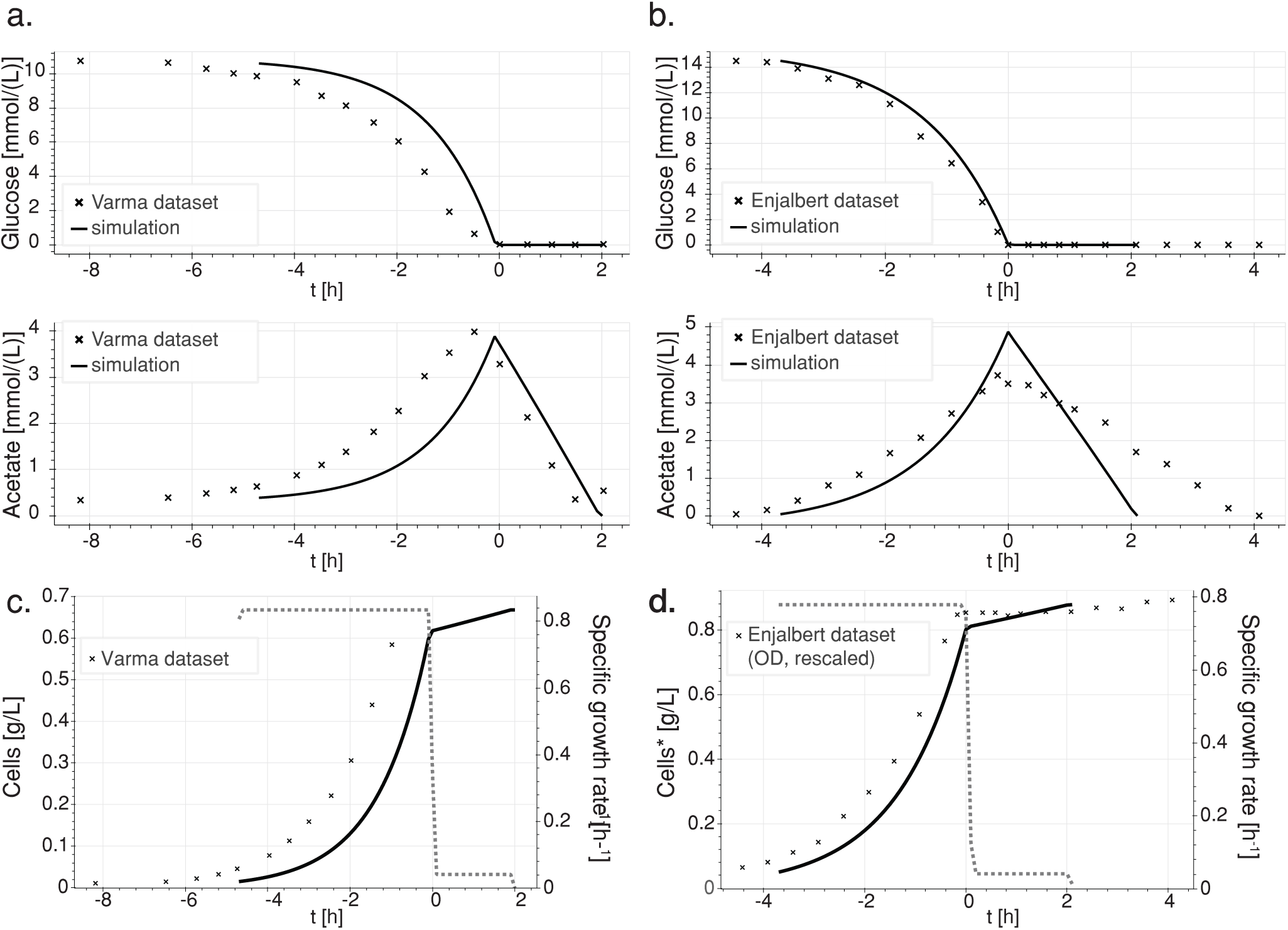
Comparison of simulated and experimental data of glucose depletion over time (Varma and Enjalbert). Simulated data represented by a solid line, experimental data by crosses. **a.** Temporal evolution of the simulated extracellular concentrations of glucose, and acetate (full lines), versus experimental data (Varma dataset) (crosses). **b.** Temporal evolution of the simulated (full lines) extracellular concentrations of glucose, and acetate (solid lines), versus experimental data (Enjalbert dateset)(crosses). **c.** Cell concentration (full line) and growth rate (dashed line) over time, simulation and Varma dataset (crosses). **d.** Cell concentration (solid line) and growth rate (dashed line) over time, simulation and Enjalbert dataset (crosses). * Experimental values were in optical density (OD600), and were linearly scaled to represent cell concentrations.

We achieved these simulated curves with no fitting. The results are the predictions of dETFL — given only the starting point of the simulation, and then aligning the curves based on the time of glucose depletion to account for experimental lag phases. The discrepancy between the simulation and the experiment data points can be attributed to several factors. First, several simulation parameters, including the maximal uptake rate for glucose, oxygen, and acetate, and the acetate maximal secretion rates, are reported with a 50% variability between the Varma and Succuro studies. We chose a common set of parameters that showed good qualitative agreement with both datasets. Changing these parameters can alter the quantitative behavior of the model, but the models always shows the same two phases. Second, variability in the experimental setup, including the *E. coli* strain, can also account for the difference in the reported glucose uptake rate by their respective authors. ME-models, and ETFL in particular, can account for the strain variability if the genetic differences (gene knock-outs, enzyme activities, enzyme over-expression) are known. Overall, these results show the dETFL framework for ME-models is able to reproduce experimental measurements of glucose uptake, acetate secretion, and biomass production in glucose-acetate diauxic growth. Our findings validate the dETFL framework as a modeling method to study the batch growth of single organisms or communities on multiple substrates and suggest its utility for investigating diauxie in mixed-substrate media.

#### Diauxic growth on glucose and lactose

Diauxic experiments show that, on a mixed medium of glucose/lactose, *E. coli* will preferentially consume glucose first, and then lactose [25, 26]. Modeling the diauxic growth of *E. coli* with dETFL should capture the lag phases and proteomic reconfiguration that are caused by the shift to a new carbon source. Therefore, this is an ideal system to challenge the ability of ME-models to describe the dynamic reorganization of the bacterial proteome. dFBA will always predict simultaneous uptake of both carbon sources, since it includes no term associated to the proteomic cost of their uptake. In contrast, ME-models describe the synthesis of enzymes, and their contribution to the overall proteome. As a result, ME-models capture the competitive allocation of the proteome to the transport of different carbon sources.

For reference, the pathways related to the glucose and lactose metabolism to G6P are summarized in Fig. 3. The figure highlights the multiple additional steps involved in the lactose pathway to form G6P, compared to the shorter glucose pathway.

**Fig 3.**
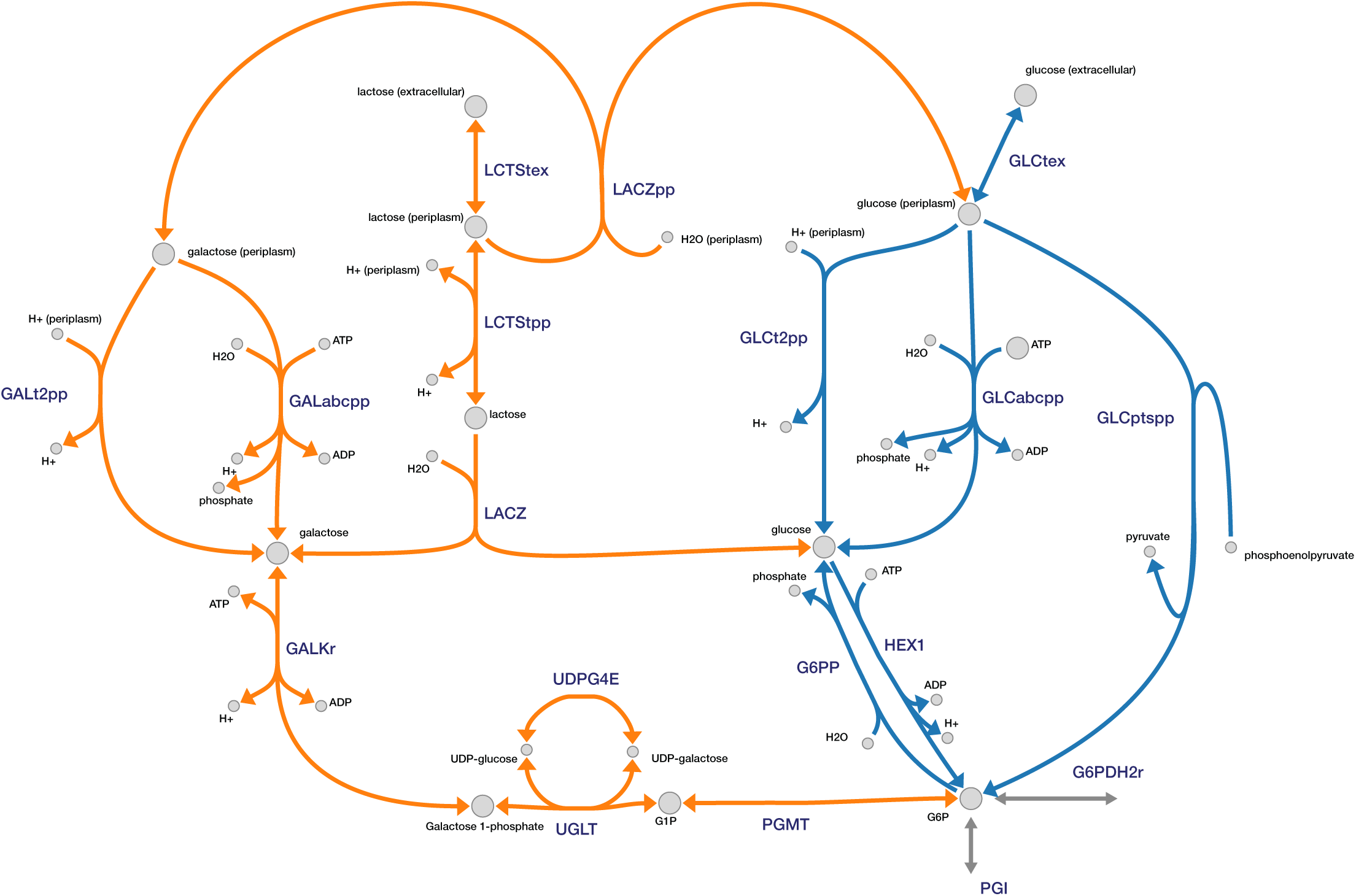
Possible uptake routes for glucose (blue) and lactose (orange), towards glucose-6-phosphate (G6P). The splitting of lactose by LACZ can be done either intracellularly, or in the periplasm. Routes towards the main central carbon metabolism are in gray. Figure made using Escher. [27]

To initialize the model for the simulation of diauxic growth, we first simulate the pre-culturing in glucose by running the model with the same standard physiology as before, with an uptake rate of 15 mmol.gDW^−1^.h^−1^ for glucose, and no lactose initially present. The model is subsequently run with these initial conditions on a mixture of glucose and lactose, at the physiologically relevant concentrations of 1 mmol.L^−1^ and mmol.L^−1^, respectively. The cell concentration is set at 0.05 g.L^−1^. After this initialization step, we ran the simulation according to the method detailed before.

The time evolution of the extracellular metabolite concentrations, cellular exchange fluxes, specific growth rate, and total biomass of the culture exhibit four phases (Fig. 4). We observe a first phase similar to the previous experiment, where glucose is taken up at a rapid rate, until its depletion, with the simultaneous production of acetate through overflow metabolism (Fig. 4-a and -b). During this phase, the growth rate is steady and high (Fig. 4-c). Relative to glucose, lactose is taken up at lower rates (Fig. 4-a). In the second phase, the specific growth rate decreases sharply while the proteome reallocates its enzymes for lactose metabolism. We also observe a drop in acetate secretion during the proteome switch and short period of acetate re-consumption. This is the lag phase, where acetate is used as a carbon source while the proteome is reconfigured to metabolize lactose. This reconfiguration shows a reduction of the total mass of enzymes that convert glucose into G6P, and an increase in the total mass of enzymes responsible for the conversion of lactose to G6P (Fig. 4-d). The third phase is characterized by a peak in lactose uptake and cell growth, followed by a decline as lactose becomes scarce. In the fourth phase, when lactose becomes scarce, the residual acetate is being taken up instead of secreted. Since it happens after lactose uptake falls below a low threshold, it indicates that lactose consumption is preferred to that of acetate. More details on the time-dependent enzyme concentrations of the glucose and Leloir pathways can be found in the Supplementary Figure S3 and S4, respectively.

**Fig 4.**
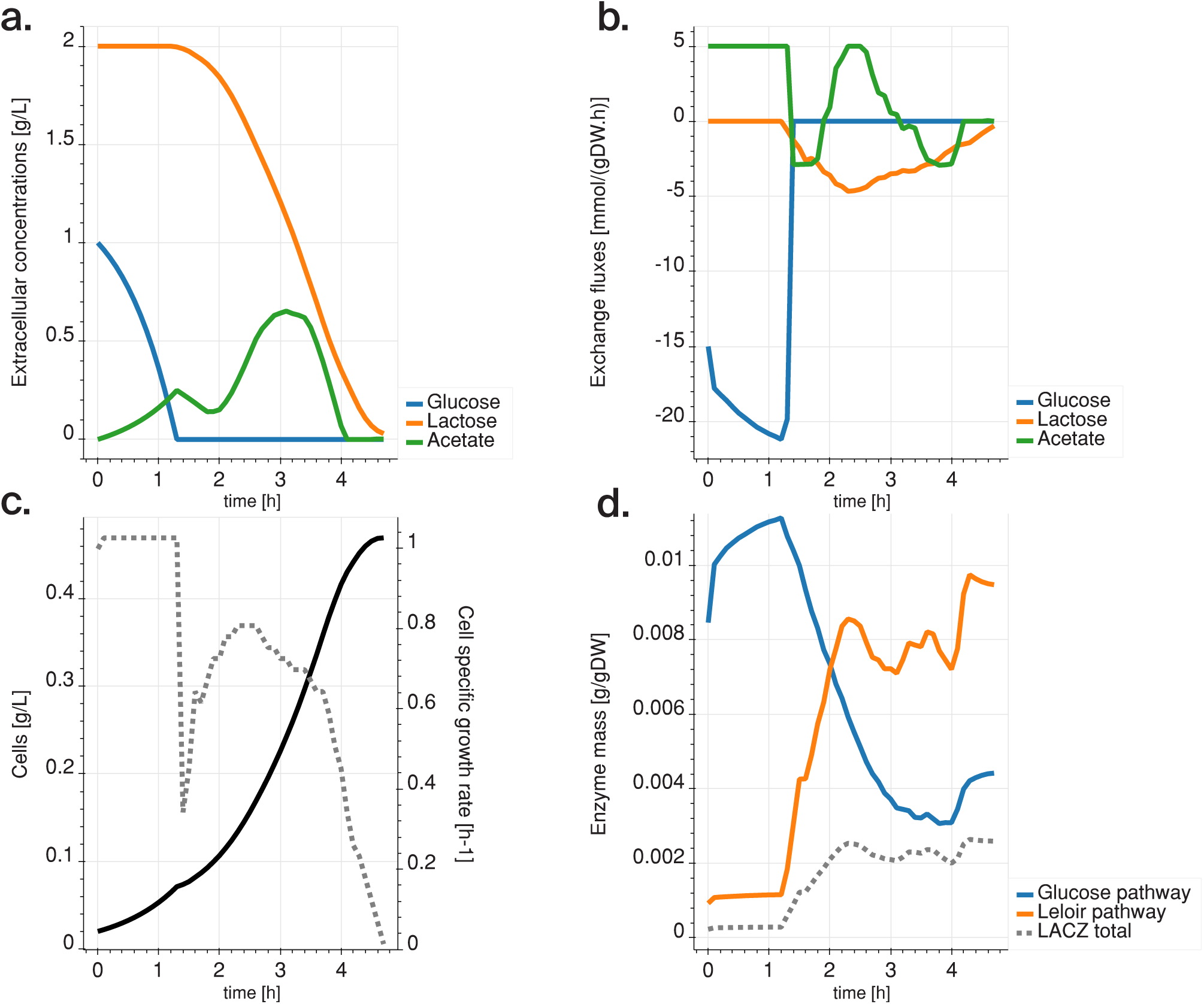
Diauxic simulation with glucose-only preculture: **a.** Temporal evolution of the extracellular concentrations of glucose (blue), lactose (orange), and acetate (green). **b.** Exchange rates of the cell. Positive exchange rates mean production, negative exchange rates mean consumption. **c.** Cell concentration (solid line) and growth rate (dashed line) of the culture over time. **d.** Mass of enzymes allocated to the transformation of glucose (blue) and lactose (orange) in G6P. The dashed gray line shows the levels of *β*-galactosidase (LACZ) enzyme (in the lactose pathway).

We next sought to assess the robustness of our diauxie prediction, and to determine whether the delayed utilization of lactose was an artifact of the initial conditions used in the simulation. We conducted a new simulation that included a preculture wherein the *E. coli* model initially only had access to lactose, with an uptake rate of 5 mmol.gDW^−1^.h^−1^.Following this preculture, we ran the simulation with identical initial conditions to previous experiments in terms of glucose, lactose, and cell concentrations.

Over the course of this simulation, we observed the same four phases : (1) preferred glucose consumption; (2) proteome switch with acetate uptake; (3) lactose consumption; and (4) acetate reutilization. Initially, glucose is taken up at a significantly smaller rate than that of the glucose preculture. The glucose uptake rate then gradually increases until glucose depletion (Fig. 5-a and -b). Comparatively, the lactose uptake rate stays low while the glucose uptake rate increases during the first phase of the experiment. The evolution of the growth rate is similar to that of the previous experiment (Fig. 5-c). Though the model was pre-cultured in lactose, the total amount of enzymes transforming lactose decreases while glucose is available (Fig. 5-d). We observe a delay, close to the cell doubling time, for initiating the utilization of glucose compared to the glucose-preculture experiment. We attribute this delay to proteome switch, from a proteome optimized for lactose consumption, to a proteome optimized for glucose consumption in this phase. In the second phase, after glucose depletion, we also observe acetate reutilization, while the enzymes needed for lactose conversion to G6P are resynthesized. In the third phase, the proteome shifts again to accommodate lactose consumption. As a result, the lactose uptake rate increases. In the final phase, acetate reutilization initiates again under scarce conditions. More details on the time-dependent enzyme concentrations of the glucose and Leloir pathways can be found in the Supplementary Figure S5 and S6, respectively.

**Fig 5.**
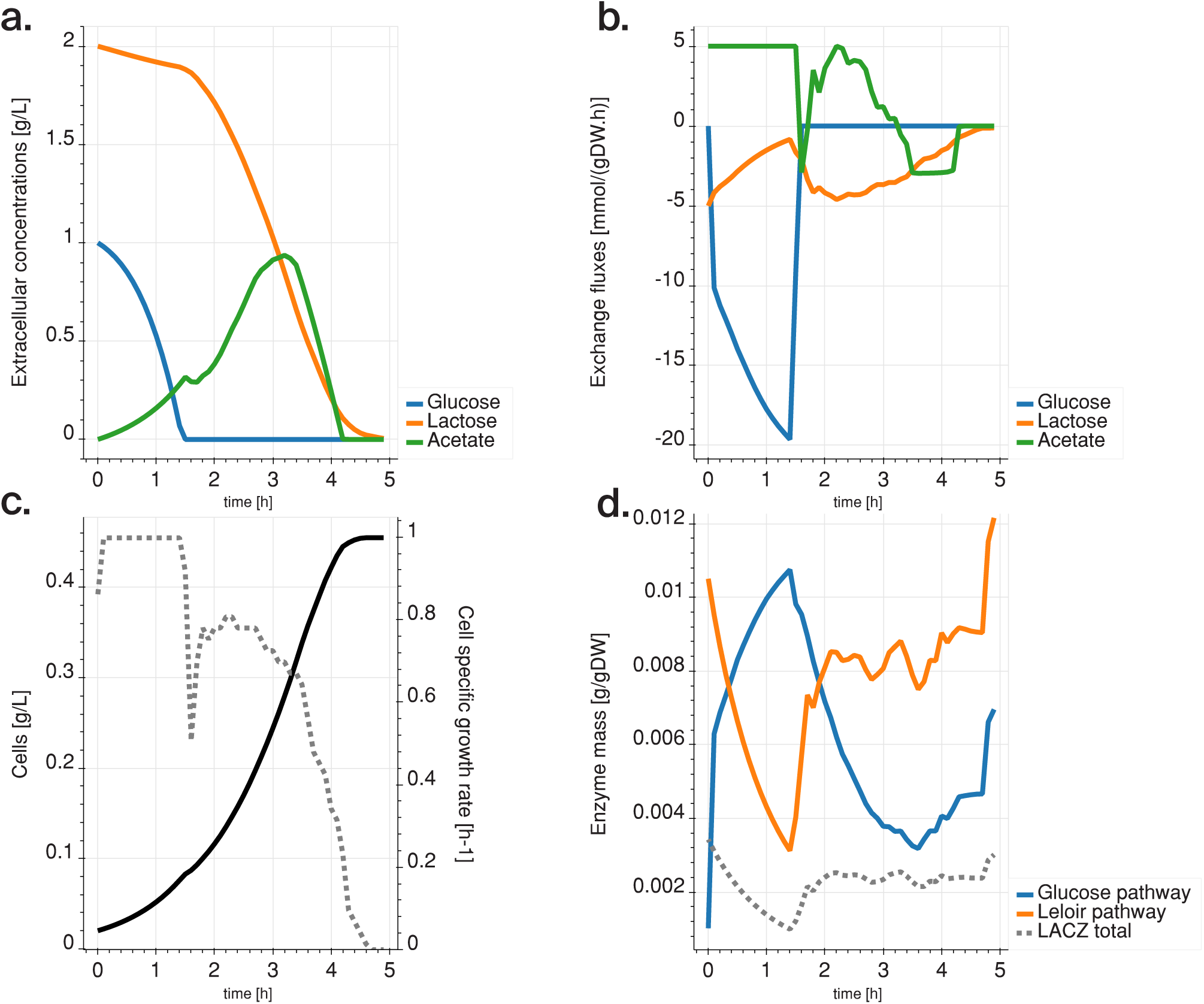
Diauxic simulation with lactose-only preculture: **a.** Temporal evolution of the extracellular concentrations of glucose (blue), lactose (orange), and acetate (green). **b.** Exchange rates of the cell. Positive exchange rates mean production, negative exchange rates mean consumption. **c.** Cell concentration (solid line) and growth rate (dashed line) of the culture over time. **d.** Mass of enzymes allocated to the transformation of glucose (blue) and lactose (orange) in G6P. The dashed gray line shows the levels of *β*-galactosidase (LACZ) enzyme (in the lactose pathway).

These simulations show strong qualitative agreement with the experimental data, for both the glucose and the lactose precultured conditions. In particular, Kremling *et al.* [26] showed a similar evolution of extracellular concentrations with a two-phase consumption of sugars. They also demonstrated that intracellular LACZ enzyme levels increase when lactose is the sole substrate left, and decrease when glucose is consumed – even after a lactose pre-culture (Fig. 4-d and Fig. 5-d). Interestingly, these agreements were achieved without adjusting any of the parameters or settings of the original ME-model. However, a key element for the consistency between the model simulations and the cellular state is a robust accounting of the intracellular states (mRNA species, enzymes and fluxes) between consecutive time steps. This has been made possible by the use of the Chebyshev centering of the cellular states in the dETFL formulation as detailed in the Methods section.

Our results strongly suggest that diauxie in *E. coli* is an optimal growth behavior. Our conceptual study suggests it is the consequence of the maximization of the cell-specific growth rate under the constraint of a limited proteome. This optimal behavior of privileging glucose consumption over lactose does not come from the pre-culturing step, but instead from the optimality of the system itself under the constraint of proteome allocation for sugar consumption. We performed additional studies and demonstrated that this behavior is not due to differences in enzyme catalytic efficiencies between the two pathways, as switching the *k*_cat_ values does not change the trend (Supplementary Figure S10). Finally, we showed that the lag time observed in experiments is determined by the proteome reallocation, and quantitavely predicted changes in the amount of enzyme for each pathway.

## Discussion

We devised both a conceptual model and a dynamic ME-model which reproduce a diauxic behavior in *E. coli*, a phenomenon that cannot be captured with current state-of-the-art models. From simulation, we determined that the preferential consumption of glucose over lactose in *E. coli* is a combined effect of its limited proteome size, enzyme properties, and substrate yield. Our model demonstrates, at the proteome level, the mechanisms of the proteome switch between conditions, and provides a method to resolve the intracellular dynamics of bacterial growth. In agreement with experimental observations, our model predicts a diauxic behavior on a medium of mixed of sugars.

In our simulations, we observed lag phases concurrent with proteome switching. The co-occurrence of the proteome reallocation and acetate reutilization suggests secreted acetate can work as an energy reserve and help the cell adapt to changing environmental conditions. The dETFL model was also able to capture different dynamic trajectories in cell fates that were dependent on the pre-culture conditions.

The preferential consumption of one carbon source vs the other is the result of an optimal trajectory of the system under the constraints of mass-balance, resource allocation and thermodynamics. These constraints are directly connected to the chemistry of the metabolic pathways in bacteria. Our conceptual model suggests that the diauxic phenomenon might be controlled through the engineering of three aspects: (i) the specific activity of enzymes (*k*_cat_), (ii) the molecular weight of the enzymes, and (iii) the number of steps involved in the substrate metabolism. The molecular weight and activity of enzymes can be altered through protein engineering, and alternative chemistries from heterologous pathways provide avenues for modifying substrate metabolism [28].

While dETFL does not account for catabolite repression, it can quantitatively describe the behavior of a cell operating under the influence of the *lac* operon. Our results imply that the genetic circuits responsible for catabolite repression are evolved as a controller to implement robust dynamic control of the optimal growth. In this regard, the catabolite repression through the *lac* operon observed in wildtype *E. coli* can be considered as a control system that ensures optimal growth of the organism. Under the selective pressure of evolution, the system might have evolved the *lac* operon to preferentially metabolize glucose in mixtures of sugars as it guaranteed an evolutionary advantage (faster growth) compared to substrate co-utilization.

As a new approach, dETFL avoids the pitfalls of simplifying modeling assumptions used in the current state-of-the-art computational models of metabolism and gene expression. Because of this, dETFL is the first dynamic ME-model formulation that can model lag phase and gradual proteome reconfiguration. However, despite these innovative findings, there are still drawbacks to dynamic constraint-based models that need refinement. For example, finding a good representative solution at each time step is extremely important. Here, we used the Chebyshev ball approach, as it is a single linear problem that is computationally simpler than other methods such as variability analysis or sampling. While we have reduced the computational burden of ME-models enough to efficiently perform iterative solving, there are new opportunities to further alleviate the computational cost of simulations. Directions to explore include fixing the integer variables of subproblems to reduce the NP-hardness of the model, and using quadratic programming, for instance, to perform an ellipsoid approximation of the enzyme solution space. Additionally, systematically reduced models, where less important parts of metabolic machinery are omitted, can also be used to reduce the complexity of the simulations [29, 30]. With a reduced computational cost of simulations, exciting new research targets are also within reach, such as the dynamic effects of gene knock-outs or drug-induced changes in cell physiology.

The new computational formulations developed herein also offer new opportunities to test other hypotheses that explain diauxie. Succurro *et al.* [24] postulated the the existence of two subpopulations of *E. coli*, where one obligately consumes glucose, while the other consumes acetate. Although the study of communities including thermodynamics-enabled ME-models is, for now, a computational challenge, cross-testing the hypothesis we present in this paper with a similar community-based context would certainly yield important insights on the respective role of proteome limitation and substrate competition in the emergence of diauxic behavior.

The inhibitory effect of glucose on certain parts of the metabolism is multiple, including catabolite repression, transient repression and inducer exclusion [31]. Moreover, more complex regulation mechanisms are found in natural environments. For example, it has been shown that, on its natural marine substate, the bacterium *Pseudoalteromonas haloplanktis* evolved regulation mechanisms allowing simultaneous diauxie and substrate co-utilization [32]. Such high-order behavior might also have its origin in an optimal growth program, and finding the biochemical constraints responsible for it would yield valuable insight on the optimal growth of organisms on complex media. In general, elucidating the emergence of regulation mechanisms in the context of evolutionary pressure will considerably increase our understanding and ability to engineer regulation systems, which are ubiquitous in biology, from wild-type *E. coli* to cancer cells. dETFL is an important step forward in this direction. Its use to uncover the optimality principles guiding the emergence of cellular regulatory control systems is key to a better understanding and, ultimately, mastery of metabolic engineering, be it applied to industrial hosts or the development of cell-based therapies.

## Supporting Information Appendix (SI)

### Supplementary Note S1

Note on the DynamicME Assumptions

### Supplementary Table S2

Properties of glucose and lactose transporting reactions and enzymes. Reaction names from the original iJO1366 model [22]. Enzyme symbols adapted from Biocyc [33]. *k*_cat_ values taken from Lloyd *et al.* [14].

### Supplementary Figure S3

Enzyme levels of the glucose pathway, in the glucose/lactose diauxie experiment with glucose pre-culture.

### Supplementary Figure S4

Enzyme levels of the lactose pathway, in the glucose/lactose diauxie experiment with glucose pre-culture.

### Supplementary Figure S5

Enzyme levels of the glucose pathway, in the glucose/lactose diauxie experiment with lactose pre-culture.

### Supplementary Figure S6

Enzyme levels of the lactose pathway, in the glucose/lactose diauxie experiment with lactose pre-culture.

### Supplementary Figure S7

2D example the different schemes to find a representative point of the space: variation analysis, sampling, or Chebyshev centering.

### Supplementary Figure S8

Dimensional example of a Chebyshev center. The feasible space is denoted by the polytope *𝒞*. The Chebyshev center with respect to variables *E*_1_ and *E*_2_ is 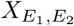. It is the center of the largest 2-D sphere on a plane parallel to (*E*_1_, *E*_2_) that is inscribed in *C*. This sphere exists on the plane *P*, materialized in light blue.

### Supplementary Figure S9

Enzyme composition of the conceptual model when no constraints are applied to the rate-of-change of enzyme concentrations.

### Supplementary Figure S10

Figure for the switched *k*_cat_ experiments.

## Materials and Methods

### Rate of change of fluxes

One of the important points in the original formulation of dFBA is that the rate at which intracellular fluxes change is constrained. In the dFBA formulation, one imposed constraint is:

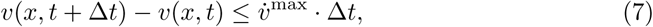

where 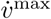 · Δ*t* is defined as the maximum change of flux between two time points. However, we the relationship between flux and enzyme concentration, as well as the dynamic mass balance, can be expressed in the following way [17]:

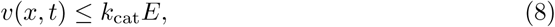

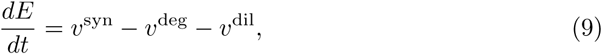

with all the rates strictly positive. From this, it directly follows that

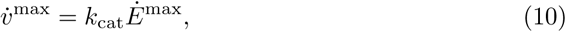

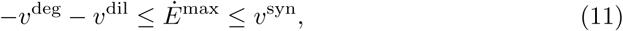

where we can rewrite 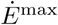 in a two components, one strictly positive, and the other strictly negative: 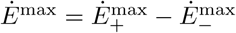. Using expression relationships from ETFL, it is hence possible to bound the maximal rate change of fluxes in a fashion that is compatible with linear programming:

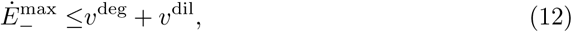

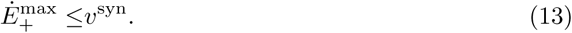

These two constraints represent, respectively, the limitation in the decrease (dilution and degradation) and increase (synthesis) of the enzyme concentration. We can rewrite these in terms of dETFL variables:

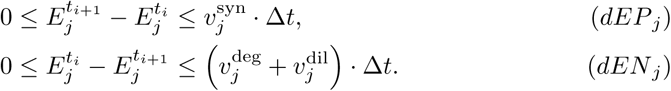

### Variability in the estimation of macromolecule concentrations

A key element in ETFL is that macromolecule concentrations are an explicit variable in the optimization problem. In dETFL, these concentrations are important because they will constraint the feasible space for the calculation of next time step.

The formulation of ETFL relies on the approximation of the growth rate of the organism by a piecewise-constant function in the dilution term of the mass balances of macromolecules. This in turn allows the linearization of the bilinear term in the mass balances. However, this approximation has an error, which is given by the resolution *η* of the discretization. Given 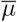 the maximum growth rate of the model, and *N* the number of discretization points, the resolution of ETFL is given by 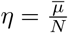. We can easily obtain the resolution of the estimation of a macromolecule concentration from this quantity.

The mass balance of a macromolecule *X* at concentration [*X*] under steady state assumption is written in ETFL:

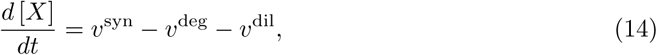

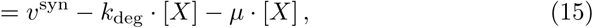

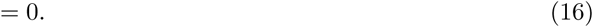

where *v*^syn^, *v*^deg^ and *v*^dil^ are respectively the synthesis, degradation and dilution rates of the macromolecule, *mu* is the growth rate, and *k*_deg_ is the degradation rate constant of the macromolecule. In ETFL, *µ* is approximated by 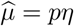, with *p ∈ {*0..*N*}. *η* is the resolution of this approximation, which means, at all times:

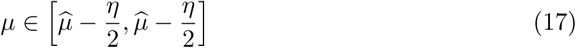

From Eq.15, and the relationship given in (17), we can rewrite:

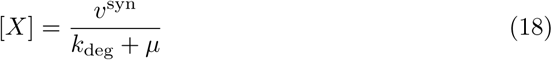

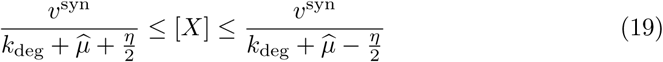

We use this expression to represent the incertitude on the macromolecules concentrations at the previous time step, which is then used to constraint the current time step.

### Backwards Euler integration scheme

At each time step, we operate an integration of the model between two time points. To this effect, using a robust integration scheme is necessary to guarantee a solution quality that is as good as possible. We chose to use a backwards (implicit) Euler integration scheme given its ability to handle stiff problems [34]. Usually, a drawback of implicit schemes is that they require to solve an implicit equation to define the state of the system at each time step. In contrast, explicit methods simply require to apply a defined set of calculations (*e.g.* a linearized state function) on the current state. In our case, however, there is little cost associated to using an implicit method rather than the explicit Euler method, since we already need to solve a whole MILP problem to compute the solution to the dETFL problem at each time step.

In this context, we can rewrite Eqs. *dEP*_*j*_ and *dEN*_*j*_ in their Euler-form:

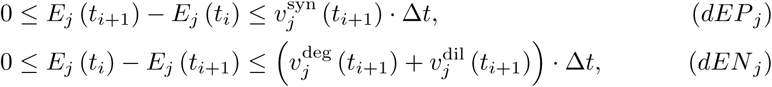

where *E*_*j*_ is the concentration of a given enzyme at the previous time step, 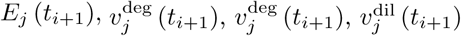 are variables of the dETFL problem in the next time step. *E*_*j*_ (*t*_*i*_) is a variable constrained around the value of the previous solution, as explained in the previous section.

### Chebyshev center

One important issue when dealing with both (mixed-integer) linear optimization and iterative solving is the multiplicity of solutions. Indeed, the optimality principle in LP only guarantees a unique global optimum value for the objective, but not a unique optimal solution for the variables. In fact, at each time point, there is most often a (piecewise-)continuum of solutions (including flux values, macromolecule concentrations …) that can satisfy a maximal growth rate, while describing different physiologies. For example, two optimal states, using different pathways with a similar enzyme cost, will yield different proteomes and associated fluxes. In addition, due to the constraints applied on the rate of change of macromolecule concentrations, in each subsequent time point, the proteome, transcriptome and flux values will be dependent on all the previous solutions. Because of these two factors, each new realisation of the integration procedure might yield different results.

An additional issue is that simplex-based solvers tend to give sparse and extremal results (corners of the explored simplex), which do not represent accurately the full extent of the considered solution space. Several methods can alleviate these issues, all based on finding a good representative of the solution space. One first solution is to use as observation the mean of the variability analysis, rather than a single optimal solution. This however requires *𝒪*(2*n*) optimizations to be carried out. Another way would be to sample the feasible space, but the sheer size of dETFL models makes sampling impractical. The Supplementary Figure S7 shows a 2-D example of the difference between these 3 approaches. The method we chose is to try to find the maximally inscribed sphere in the solution space. The center of this sphere, called the Chebyshev center, can be found by optimising a single linear problem if the solution space is a polytope [35]. It is the case in dETFL, as the problem is defined with linear inequality constraints.

In the case of a polyhedron defined by inequalities of the form 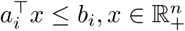, finding the Chebyshev center of the solution space amounts to solving the following optimization problem:

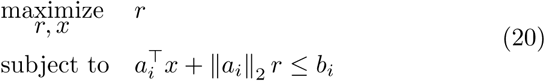

This is similar to adding a common slack to all inequalities and maximize the its size, which maximizes the distance of the solution to the inequality constraints. However, not all variables and constraints need to be considered in the definition and inscribing of this sphere. In particular, we are interested in a representative solution for macromolecule concentrations, which only play a role in a limited set of constraints. To this effect, we define ℐ_*c*_ and 𝒥_*c*_, respectively the set of inequality constraints and variables with respect to which the Chebyshev center will be calculated. Let us also denote *ℰ* the set of equality constraints of the problem, *a*_*i*_, *c*_*i*_ respectively the left-hand side of the inequality and equality constraints, and *b*_*i*_, *d*_*i*_ their respective right-hand side. From there, we can define the modified centering problem:

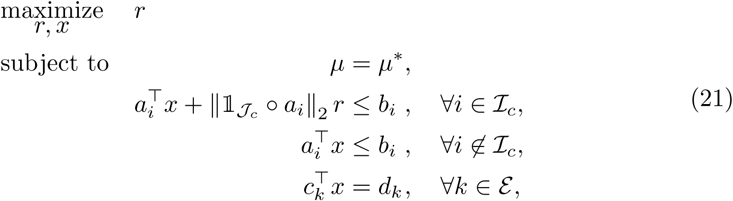

where *µ\** is the maximal growth rate calculated at this time step, *r* the radius of the Chebyshev ball, *x* the column vector of all the other variables of the ETFL problem, 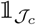 has for *j*^th^ element 0 if *j ∈ 𝒥*_*c*_, else 1, and ∘ denotes the element-wise product between two vectors. Thus, 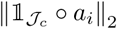 is the norm of the projection of the constraint vector onto *𝒥*_*c*_. We show an example illustration in 3D in the Supplementary Figure S8.

For enzymes, for example, it is akin to making the model produce more enzymes than necessary to carry the fluxes, while respecting the total proteome constraint. By maximizing the radius of the sphere inscribed in the solution space, at maximal growth rate, we are effectively choosing a representative solution of the maximal growth rate feasible space. We then use this solution as a reference point for the next computation step.

All simulations in this paper perform Chebyshev centering on enzyme variables at each time step.

### Initial conditions

Since dETFL is an iterative method, it is necessary to set an initial reference point (initial conditions) from which the dynamic analysis will integrate over time. The initial solution is set up as follows:

1. Set typical uptake fluxes for carbon sources and oxygen,
2. Perform a growth maximization using ETFL
3. Fix the growth to the optimum,
4. Find the Chebyshev center of the solution space.

The solution reported by the latter optimization problem is then used as a starting solution for the dETFL analysis.

### Extracellular concentrations

At each time step, extracellular concentration are updated following a standard Euler scheme, similarly to what is done in Mahadevan *et al.* [13]. The extracellular concentrations of glucose, lactose, and acetate, follow a system of ordinary differential equations:

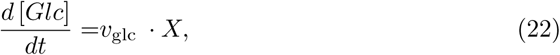

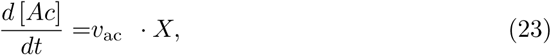

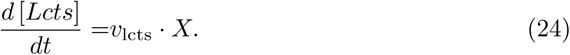

We linearize this system into the following forward Euler scheme:

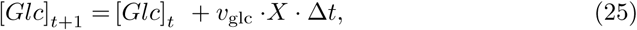

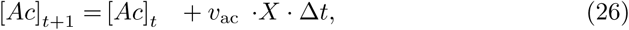

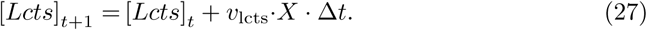

We use these linearized equations to update the extracellular medium after the solution to each time step has been computed.

### Model

The model used is the the vETFL model of iJO1366, presented in the original ETFL publication [17]. 15 additional enzymes were added to the model to properly account for the protein cost of transporting glucose, lactose and galactose from periplasm to the cytoplasm. A simplified metabolic map of the glucose, lactose, and galactose pathways to G6P is shown in Fig. 3.

### Kinetic information

The Michaelis-Menten parameter 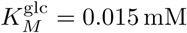 for glucose was taken from the original dFBA paper [13]. The 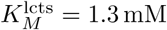 of for lactose was obtained from a study by Olsen *et al.* on the specificity of lactose permeases [36]. Details on the added enzymes are available in the Supplementary Table S2. The Michaelis-Menten parameter 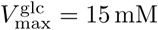 was used similarly to previous work [13].

Because of incertitude in the values found in the literature, 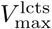 was directly computed from the catalytic rate constants of enzymes consuming periplasmic lactose (LACZpp, LCTStpp, LCTS3ipp). Since ETFL gives access to enzyme concentrations, we can rewrite the expression of 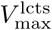 using catalytic rate constants 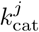:

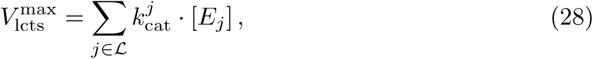

where ℒ is the set of periplasmic enzymes consuming lactose. Taking this into account allows to replace the parameter 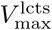 by an explicit internal variable.

Acetate transport is assumed to be mostly diffusive [37], and its secretion rate was bounded at 5 mmol.gDW^−1^.h^−1^, and its uptake to 3 mmol.gDW^−1^.h^−1^. Oxygen is assumed to be non-limiting, and is given a maximal uptake rate of 15 mmol.gDW^−1^.h^−1^. These values are of the same order of magnitude as in previous studies [5, 13, 24]

### Implementation

The code has been implemented as a plug-in to pyTFA [38], a Python implementation of the TFA method, and ETFL [17], an implementation of ME-Models accounting for expression, resource allocation, and thermodynamics. It uses COBRApy [39] and Optlang [40] as a backend to ensure compatibility with several open-source (GLPK, scipy) as well as commercial (CPLEX, Gurobi) solvers. The code is freely available under the APACHE 2.0 license at https://github.com/EPFL-LCSB/etfl.

## Acknowledgements

The authors would like to thank Dr. Ljubiša Mišković and Dr. María Masid Barcón for valuable discussions around this project; and Dr. Kaycie Butler for her valuable input on the wording and flow of this manuscript. Pierre Salvy would like to express his gratitude to Kilian Schindler and Prof. Daniel Kuhn for the valuable discussion around the formulation of the Chebyshev problem. This work has received funding from the European Union’s Horizon 2020 Research and Innovation Programme under the Marie Sklodowska-Curie grant agreement No 722287, the European Union’s Horizon 2020 Research and Innovation Programme under grant agreement No 686070, and the Ecole Polytechnique Fédérale de Lausanne (EPFL).

