## Supplementary material for "Emergence of diauxie as an optimal growth strategy under resource allocation constraints in cellular metabolism": SI1-10

Salvy et al.

#### Supplementary note S1: Discussion on the assumptions in DynamicME

Lloyd *et al.* [1] developed an efficient ME-model for *E. coli*, and Yang *et al.* used it to formulate a dynamic analysis framework (dynamic-ME) [2] similar to dynamic flux balance analysis (dFBA) [3]. However, in order to handle the computational complexity of their model, they introduced a number of assumptions which resulted in several limitations to their model. In particular, the following items might be limiting:

- The standard solving procedure uses a dedicated quad-precision solver [4] and its assorted solving algorithm [5].
- Moreover, an important assumption in this method is that the dynamic algorithm approximates uptake fluxes bounds to be constant as long as the substrate in question is not depleted, which is neglects the impact of kinetic laws at different substrate concentrations.
- It also assumes the proteome does not change during that time.
- Additionally, the efficiency of their formulation comes from the use of equality constraints between the metabolic fluxes and the catalytic availability of the enzymes.
- As a result, these models cannot predict the presence of enzymes that do not carry flux, as can be the case in a transition phase between two phenotypes.
- More importantly, this method does not tackle the problem of alternative solutions at a given time-step, and hence does not acknowledge the possibility of different time traces depending on the solution choice.
- Finally, this method does not allow modeling thermodynamics constraints.

#### Supplementary Table S2

| Reaction | Reaction name | Enzyme Symbol | $k_{cat}$ [$s^{-1}$] |
|---|---|---|---|
| GLCabcpp | Glucose transport via the ABC system | GLCt2pp_ABC_18 | 120.0 |
| GLCt2pp | Glucose transport via proton symport | GLCt2pp_GALP | 40.1 |
| GLCptspp | Glucose transport via PEP to Pyruvate PTS | GLCptspp_157 | 134.5 |
| | | GLCptspp_164 | 139.2 |
| | | GLCptspp_165 | 135.3 |
| HEX1 | Hexokinase (glucose:ATP) | HEX1_GLUCOKIN | 279 |
| LACZpp | $\beta$-galactosidase (periplasmic) | LACZpp_EG12013 | 58.1 |
| LACZ | $\beta$-galactosidase (cytoplasmic) | LACZ_BETAGALACTOSID | 211 |
| LCTStpp | Lactose transport via proton symport | LCTStpp_LACY | 37.5 |
| | | LCTSt3ipp_YDEA | 35.0 |
| | | LCTSt3ipp_B0070 | 35.1 |
| | | LCTSt3ipp_B2170 | 35.2 |
| GALKr | Galactokinase | GALKr_G7096 | 38.0 |
| | | GALKr_GALACTOKIN | 34.3 |
| UGLT | UDPglucose–hexose-1-phosphate uridylyltransferase | UGLT_GALACTURIDYLYLTRANS | 62.0 |
| UDPG4E | UDPglucose 4-epimerase | UDPG4E_UDPGLUCEPIM | 128 |
| GALabcpp | Galactose transport via the ABC system | GALabcpp_ABC_18 | 120.0 |
| | | GALabcpp_ABC_46 | 110.0 |
| GALt2pp | Galactose transport via proton symport | GALt2pp_GALP | 40.1 |

Table S1: Properties of glucose and lactose transporting reactions and enzymes. Reaction names from the original iJO1366 model [6]. Enzyme symbols adapted from Biocyc [7]. $k_{cat}$ values taken from Lloyd *et al.* [1].

### Supplementary Figures S3-6: enzyme levels of pathways depending on the preculture conditions

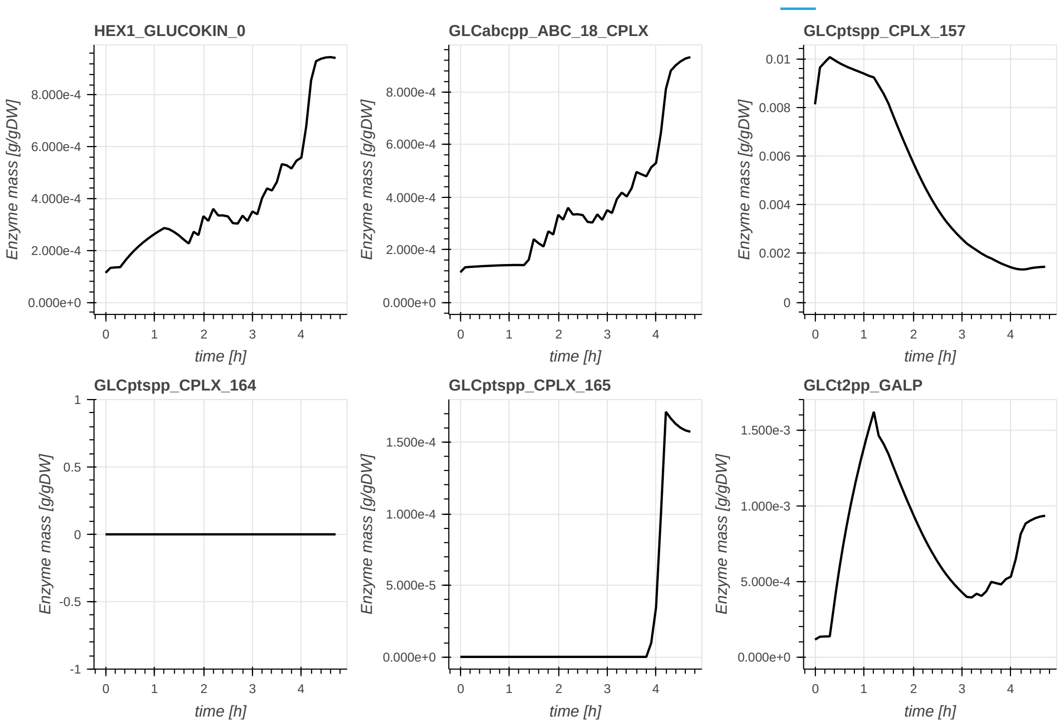


Figure S3: Enzyme levels of the glucose pathway, in the glucose/lactose diauxie experiment with glucose pre-culture.

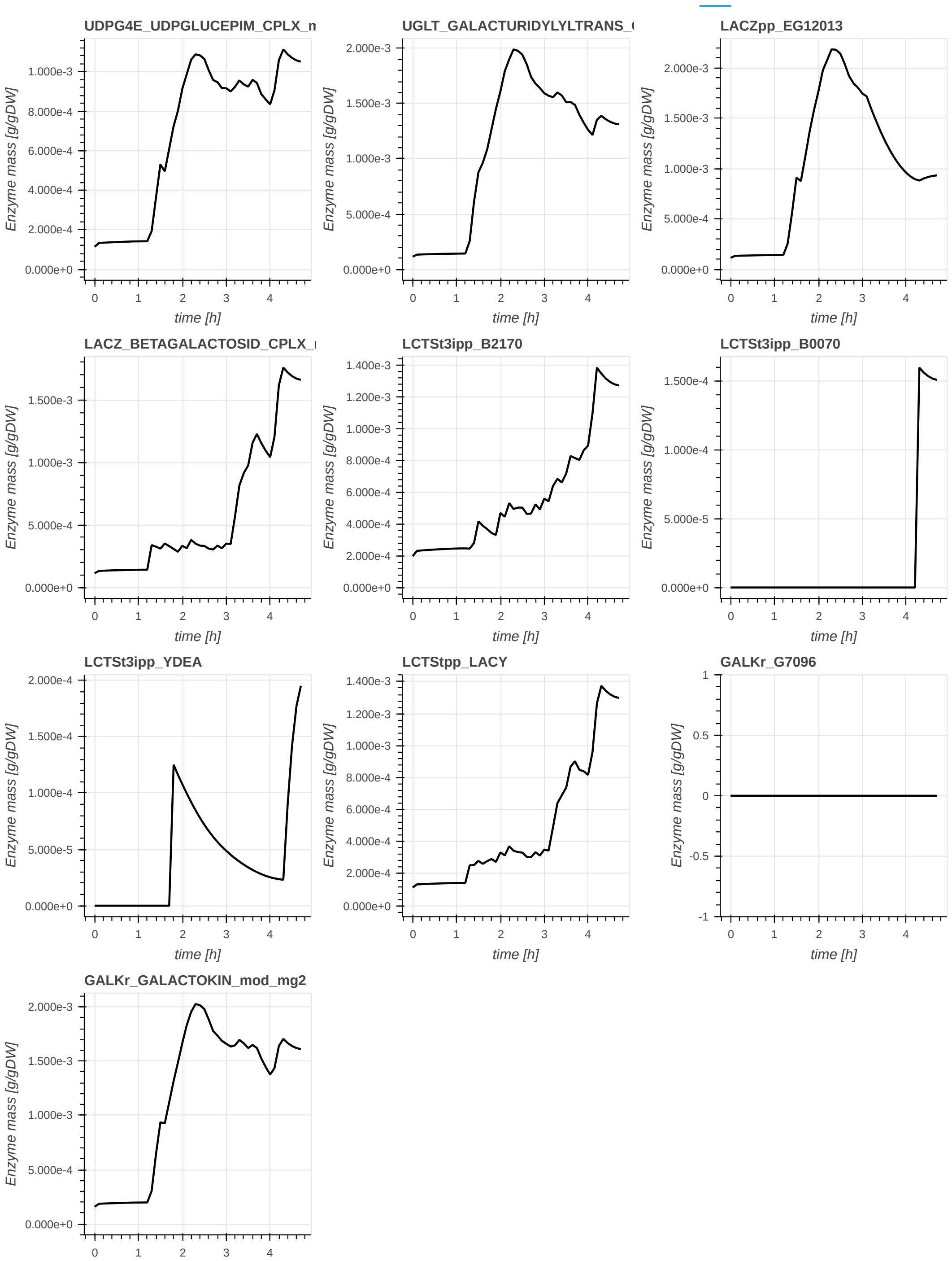


Figure S4: Enzyme levels of the lactose pathway, in the glucose/lactose diauxie experiment with glucose pre-culture.

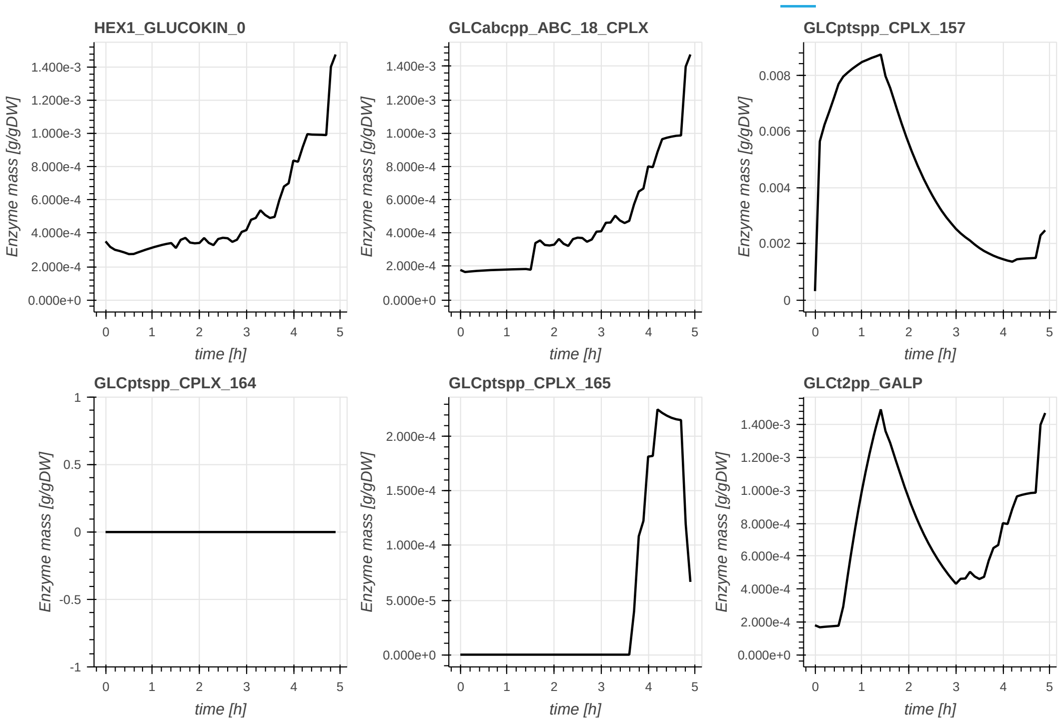


Figure S5: Enzyme levels of the glucose pathway, in the glucose/lactose diauxie experiment with lactose pre-culture.

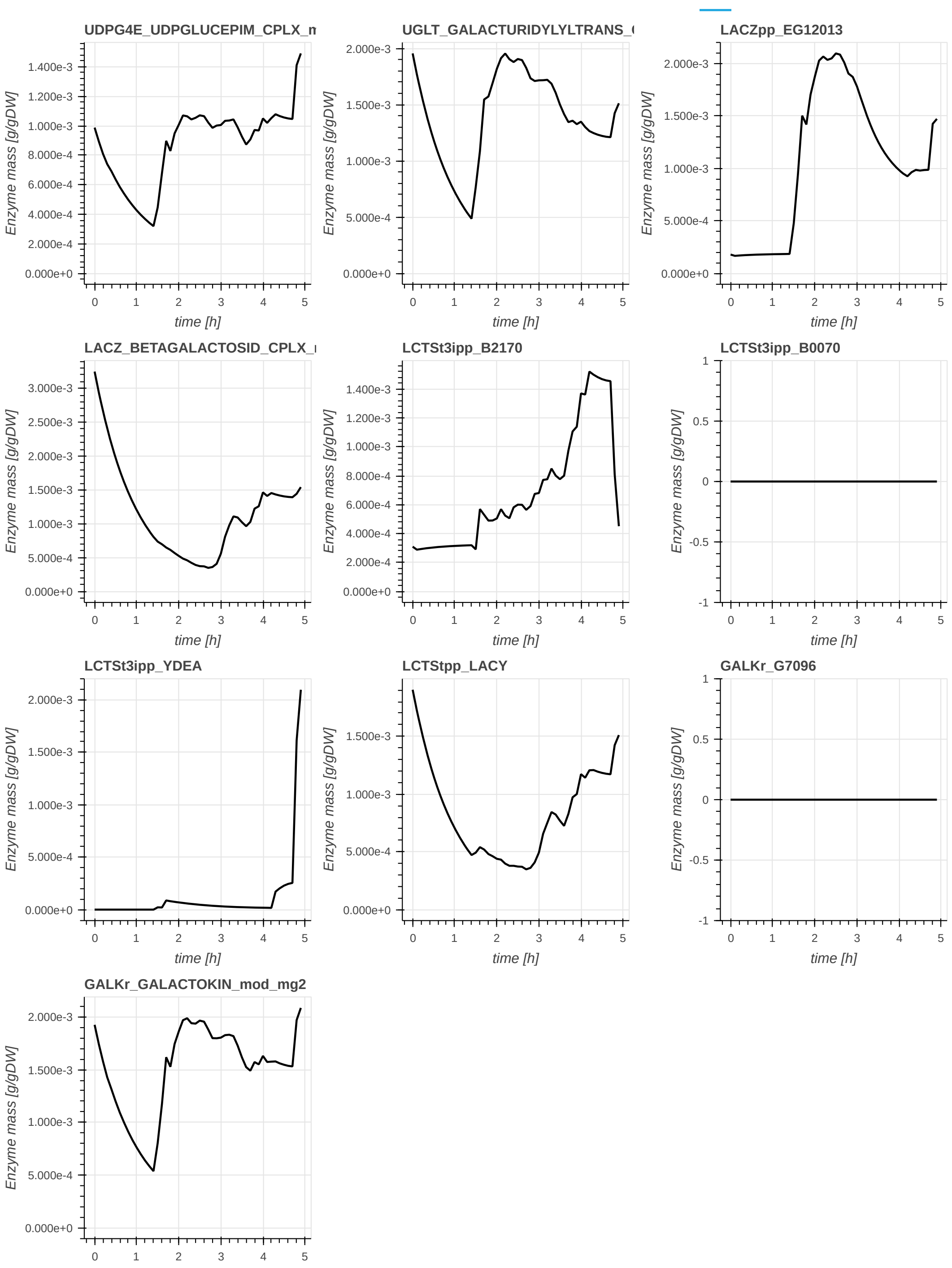


Figure S6: Enzyme levels of the lactose pathway, in the glucose/lactose diauxie experiment with lactose pre-culture.

#### Supplementary Figures S7-8: Chebyshev centering

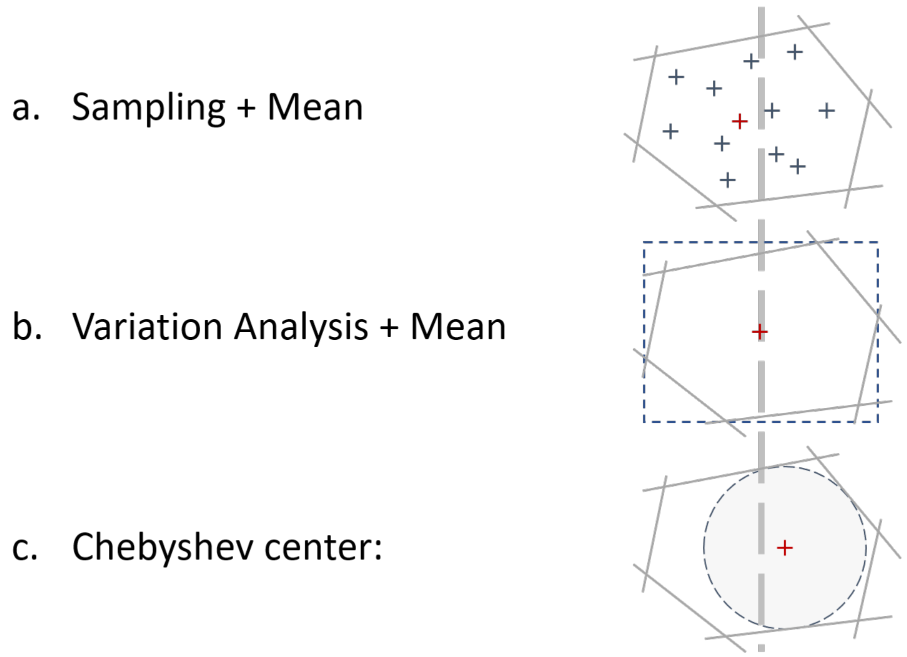


Figure S7: 2-Dimensional representation of different schemes to represent the solution space (gray polygon). The methods do not yield similar results (gray dashed line through the figures). **a.** It is possible to sample solutions (blue crosses) within the solution space, and take their mean (red cross) as a representative solution. The mean is still part of the solution space due to the convexity of the problem. **b.** Variation analysis (successive minimization/maximization of variables) allows to find the minimal bounding box (blue dashed lines) around the solution space. The center of the box (red cross) is also in the solution space, and can be used as a representative solution. **c.** The Chebyshev method finds the largest topological ball (grayed area) that can fit in the solution space. The center of the ball, or Chebyshev center (red cross) is also part of the solution space and can be used as a representative solution.

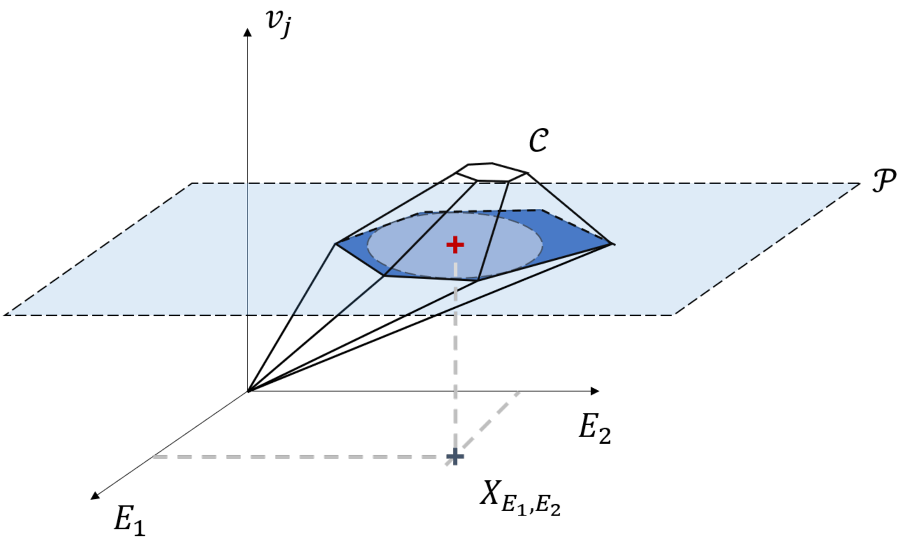


Figure S8: 3-Dimensional $(E_1, E_2, v_j)$ example of a Chebyshev center. The feasible space is denoted by the polytope $\mathcal{C}$. The Chebyshev center with respect to variables $E_1$ and $E_2$ is $X_{E_1,E_2}$. It is the center of the largest 2-D sphere on a plane parallel to $(E_1, E_2)$ that is inscribed in $\mathcal{C}$. This sphere exists on the plane $\mathcal{P}$, materialized in light blue.

#### Supplementary figure S9: Enzyme composition of the conceptual model when no constraints are applied to the rate-of-change of enzyme concentrations

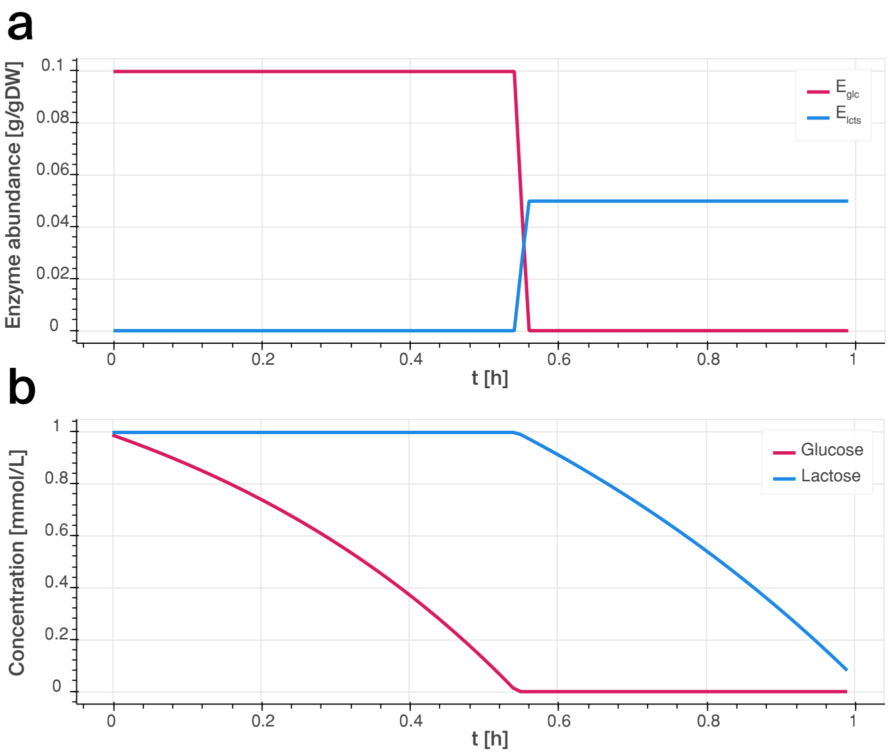


Figure S9: Enzyme composition of the conceptual model when no constraints are applied to the rate-of-change of enzyme concentrations. **a.** Enzyme content over time for the conceptual model on a mixed substrate. Glucose enzymes in pink, lactose enzymes in blue. **b.** Content of the batch reactor over time: Glucose (pink), lactose (blue).

#### Supplementary figure S10: dETFL results with switched $k_{cat}$ between the glucose and Leloir pathways

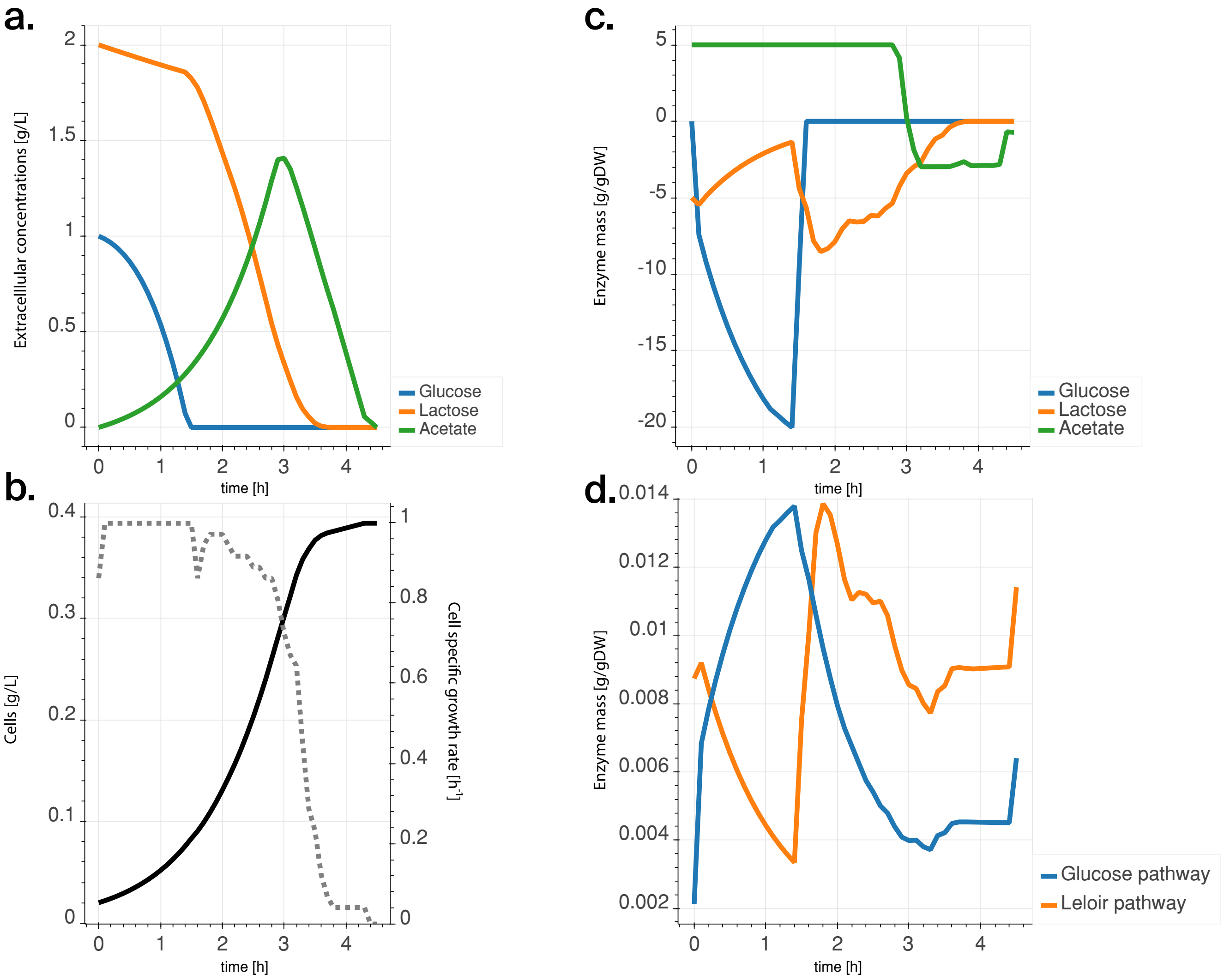


Figure S10: Results of the diauxic simulation with lactose-only preculture, and switched $k_{cat}$ values between the glucose and Leloir pathways (resp. $37.7\,s^{-1}$ and $135\,s^{-1}$: **a.** Temporal evolution of the extracellular concentrations of glucose (blue), lactose (orange), and acetate (green). **b.** Cell concentration (full line) and growth rate (dashed line) of the culture over time. **c.** Exchange rates of the cell, same colors as in subfigure -a. Positive exchange rates mean production, negative exchange rates mean consumption. **d.** Mass of enzymes allocated to the transformation of glucose (blue) and lactose (orange) in G6P. The dashed gray line shows the levels of $\beta$-galactosidase (LACZ) enzyme (in the Leloir pathway).
